# Exploiting homologous recombination increases SATAY efficiency for loss- and gain-of-function screening

**DOI:** 10.1101/866483

**Authors:** Agnès H. Michel, Sabine van Schie, Andreas Mosbach, Gabriel Scalliet, Benoît Kornmann

## Abstract

The analysis of large-scale transposon mutant libraries is becoming a method of choice for functional genomics in bacteria and fungi. We previously established SAturated Transposon Analysis in Yeast (SATAY) to uncover genes necessary for growth in any condition in *S. cerevisiae* (Michel et al., 2017). We present an improved version leveraging homologous recombination to increase transposition efficiency by a factor 10, allowing a single experimenter to rapidly perform several parallel screens. We demonstrate its potential by presenting (1) a comparison of the essential gene sets between two yeast laboratory backgrounds, (2) a comprehensive description of essential genes displaying phenotypic delays – we highlight their common features and propose plausible explanations for this phenomenon –, (3) a genome-wide analysis of loss- and gain-of-function mutations conferring sensitivity or resistance to a compendium of 9 anti-fungal compounds. This study highlights the power of this improved SATAY protocol for yeast functional- and pharmaco-genomics.

## Introduction

The yeast *Saccharomyces cerevisiae* continues to be an instrumental model system to understand the underpinnings of eukaryotic life. Several methods allow genomewide screening of mutant phenotypes to pinpoint and catalogue genes involved in any cellular function (Weissman et al., 2010). These methods usually include the ordered arraying or pooled growth of deletion libraries (Giaever et al., 2002). In the latter case, each deletion strain bears a distinctive DNA “barcode” for subsequent identification. Recent years have witnessed an explosion of research exploiting these tools (Giaever and Nislow, 2014).

There are, however, limitations to the use of deletion libraries. Indeed, as generating libraries is extremely labor-intensive, the set of independent libraries available remains narrow and is used repeatedly, confining screens to few genetic backgrounds. Because genetic background significantly affects the penetrance and/or expression of genetic traits, this complicates the connection from genotype to phenotype. Moreover, the repeated amplification of deletion libraries is associated with the emergence of genetic adaptations. (Teng et al., 2013) for instance estimated that 56% percent of the strains in a commonly used deletion library bear secondary mutations.

Recently, an alternative approach has been pioneered to interrogate the genome of a host of microorganisms, including *S. cerevisiae*. This approach utilizes dense libraries of transposon mutants that are easily generated as needed, in a background of interest. Libraries are grown as pools, and instead of using barcodes, mutations are directly identified by next-generation sequencing of the transposon-genome junctions (Christen et al., 2014; Coradetti et al., 2018; Edskes et al., 2018; Girgis et al., 2007; Guo et al.,2013; Sanchez et al., 2019; Segal et al., 2018; Uhse et al., 2018; van Opijnen et al., 2009; Zhu et al., 2018). We have recently developed a protocol dubbed SATAY (for SAturated Transposon Analysis in Yeast), which utilizes a galactose-inducible *Ac* transposase to mobilize a minimal transposon (mini*Ds*) (Michel et al., 2017). Mini*Ds* is originally located within, and interrupts, the *ADE2* gene (Weil and Kunze, 2000). Excision of the mini*Ds* element causes the two halves of the *ADE2* genes to be ligated by non-homologous end-joining (NHEJ) (Lazarow et al., 2012), thereby reconstituting an active gene and conferring adenine prototrophy. By inducing transposition on 200-300 +galactose -adenine standard Petri dishes, this protocol generates 1.5-3 millions of transposon mutants, of which ~300,000-500,000 independent transposon insertions are routinely detected. The density of transposon insertions makes it possible to identify regions in which insertions affect growth, either positively or negatively, in any given set of conditions. It also allows a sub-ORF resolution where transposon insertions in different parts of an ORF cause truncation of functional domains, yielding loss-, gain- and separation-of-function alleles (Michel et al., 2017).

Although this protocol requires no specific infrastructure or equipment, the manpower involved in pouring, plating and harvesting 200 to 300 plates limits the throughput attainable by a single research group.

Here we show that the incorporation of a short “repair template” allowing the reconstitution of *ADE2* via accurate homology-directed repair, instead of imprecise NHEJ, increases the efficiency of the protocol by 10-fold. We also develop a fully “liquid” protocol, which decouples the transposition from the selection of the transposed mutants and bypasses the need for solid medium. We show that these screens can be multiplexed and, as a proof-of-principle, we investigate the effect of various well-characterized bioactive and antifungal compounds on the growth of these mutants.

## Results and Discussion

### Improving the SATAY protocol with DNA repair templates

The original SATAY protocol yields 8-11.10^3^ colonies per 8.5- Ø cm Petri dish from ~10^8^ cells plated on transposition inducing medium, i.e. a ratio of 1/10,000. Plating and harvesting ~300 Petri dishes is necessary to reach a resolution of one insertion per 10-40 bp. These steps are thus rate limiting in the SATAY approach. Furthermore, background-specific variations in transposition efficiency hinders the amenability of the approach in some laboratory strains. Of particular interest is the W303 background, widely used in studies on DNA repair (Thomas and Rothstein, 1989), in which systematic deletion/overexpression libraries for high-content screening are not available. Compared to BY4741 (the background in which the SATAY approach was originally established), W303 transposed with a ~10-fold reduced efficiency. Reaching saturation in W303 would thus have required several thousand plates.

This prompted us to investigate the factor(s) limiting transposition efficiency. One obvious candidate was the availability of the transposase, which was ruled out as expressing the transposase from a multicopy 2μ episomal vector did not increase the transposition rate, but instead decreased it by at least 30-fold.

We then reasoned that a limiting step could be the repair of the *ADE2* gene necessary for the generation of Adenine prototrophic colonies. In the original SATAY protocol, the excision of the mini*Ds* transposon by the transposase must be followed by non-homologous joining of the two free DNA ends to reconstitute a functional *ADE2* gene. As observed previously (Lazarow et al., 2012), this step is error-prone and leaves “scars” of various length at the joining point, which might cause a substantial fraction of the repair events to lead to a non-functional *ADE2* gene (Fig. 1D, left). Moreover, *S. cerevisiae* is notoriously inefficient at NHEJ (Boulton and Jackson, 1996).

**Fig. 1.**
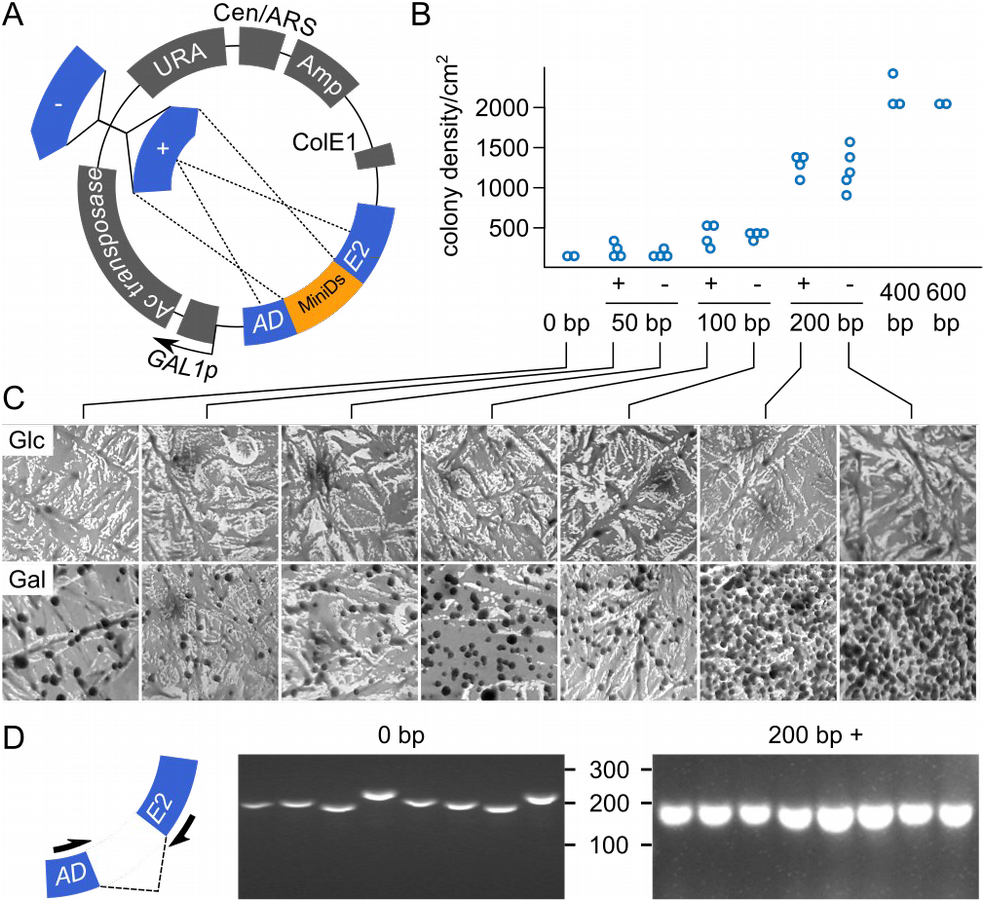
Design of plasmids with repair templates. A) Plasmid maps. The repair templates, homologous to the part of ADE2 that is interrupted by the transposon, are shown as blue arrows labeled (+) or (–), depending on their orientation. B) Number of Ade+ clones obtained on -adenine +galactose medium for the various constructs containing no repair template (pBK257, original SATAY construct), or repair templates of the indicated orientation and length. C) Representative pictures of the surfaces of plates in transposase non-inducing (Glc) and inducing (Gal) conditions after 10 days. D) PCR of “scars” around miniDs original position, after excision of the transposon in the absence (left) or presence (right) of a 200-bp repair template.

We thus sought to “help” the repair of the *ADE2* gene by incorporating templates for homology-directed repair (HDR) of the excision site into the original construct (Fig. 1A). Because HDR crucially depends on the length of homology regions (Hua et al., 1997), we initially tested three different lengths (50, 100 and 200 bp) in both possible orientations (+ and -). We plated ~ 10^8^ cells onto +galactose -adenine medium, according to the original protocol, and monitored the appearance of Ade+ clones. The introduction of a repair template increased transposition efficiency up to 7-fold (Fig. 1B, C). By contrast, the orientation of the template had no apparent effect. To assess whether *ADE2* repair was happening via HDR rather than NHEJ, we PCR-amplified the “scar” region within *ADE2* around the original location of mini*Ds* from several Ade+ clones. Contrary to the length heterogeneity, typical of NHEJ, observed upon repair without repair template (Fig. 1D, left), scar regions from plasmids bearing a 200-bp repair template were of homogeneous length (Fig. 1D, right), indicating that the repair had taken place by HDR.

The increased efficacy with increased template length prompted us to test two additional constructs with 400 and 600 bp repair templates. Because orientation had negligible effects, we pursued only with plasmids in the (+) orientation. The 400-bp template yielded more clones than the 200-bp ones. The 600-bp template did not noticeably improve transposition over the 400-bp one, indicating that the length of the repair template did not influence repair beyond 400-600 bp (Fig. 1B). We therefore continued with the 600-bp construct.

We estimated that the presence of the 600 bp repair template boosted the transposition by ~10-15 fold compared to the previous NHEJ-only design.

With this new protocol we generated two libraries in the common laboratory strains BY4741 and W303.

### A liquid SATAY protocol

We wondered whether we could leverage the increased efficacy to simplify the SATAY protocol substantially. In the original protocol, a strain is plated densely on +galactose -adenine plates and allowed to transpose for three weeks. With 8-11·10^3^ colonies per plate, ~300 plates were needed to obtain 2-3·10^6^ independent clones. This long step was necessary to reach the highest possible number of clones. It was carried out on solid medium to allow late-emerging clones to catch-up with the early emerging ones, and therefore to minimize variability arising from stochastic differences in transposition timing. With the increased efficiency, it became likely that the long waiting step on solid medium was no longer required, and that transposition could happen in liquid culture (Fig. 2A). To test this idea, we induced a culture of BY4741-derived yeasts harboring the new construct in -uracil (to select for the plasmid) +galactose medium at an original OD of 0.2, and plated aliquots of the culture at different time points. After an initial period where the culture grew but the number of Ade+ colonies increased only modestly, the culture entered a saturation phase, during which the number of new Ade+ colonies increased sharply. After two days, a plateau was reached with 6·10^4^ Ade+ cells per ml, i.e. ~1/1000 of the total (Fig. 2B). Because more than 80% of Ade+ cells appeared during the saturated phase, they likely arose from independent transposition events. Thus a comprehensive SATAY library of 2-3·10^6^ distinct clones could be obtained in two days with as little as 30-50 ml of culture.

**Fig. 2:**
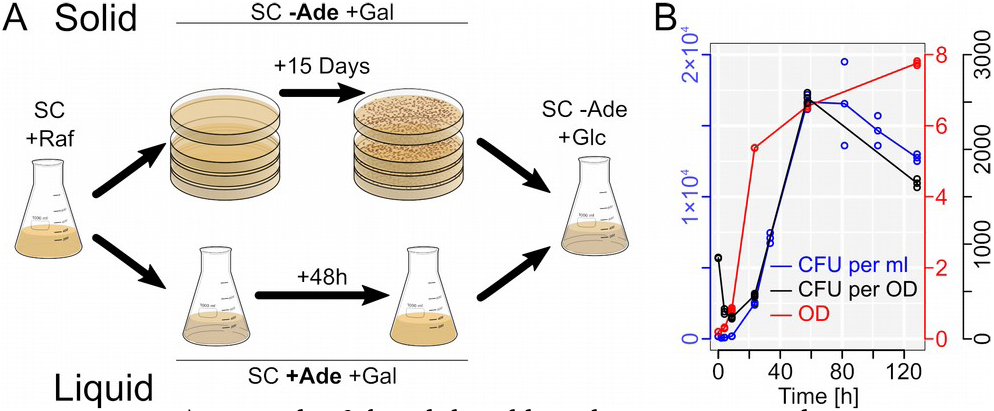
A) Principle of the solid and liquid SATAY protocols. B) Appearance of Ade+ colony forming units (CFU) upon transposition induction in SC -uracil +galactose. Blue, Ade+ CFU per ml culture; red, Optical density (OD) of the culture; black, number of Ade+ CFU per OD.

We thus set out to reproduce this in larger scale in both BY4741 and W303 backgrounds. After 52 to 66 hours of induction in -uracil +galactose medium, we obtained 1.5-2·10^4^ Ade+ cells/ml in the By4741 background (in three independent libraries) and ~20·10^4^ Ade+ cells/ml in the W303 background (one library). These numbers were inferred by retrospective counting (see materials and methods). We selected for transposed clones by inoculating a total of 260 ml in 8 L of -adenine +glucose medium for the BY4741 library, and 60 ml in 2 L for W303. We thus seeded 4·1ø^6^ and 13·10^6^ independent transposed clones at an initial OD of 0.2 for By4741 and W303, respectively. After ~70 hours, the OD had reached ~1-4. We then harvested the cultures, measured the ratio of Ade+ vs Ade-cells and found it to be 45-70%. We isolated genomic DNA and proceeded with the standard SATAY procedure as previously described (Michel et al., 2017).

Transposon insertion data for these and previous libraries can be browsed here: http://genome-euro.ucsc.edu/s/benjou/Lsz2Gbt4

### Comparison of the BY4741 and W303 backgrounds

We compared the two libraries generated in W303 (solid and liquid protocols) to libraries generated in BY4741 (solid and liquid protocols from this study plus seven previously published libraries, Michel et al., 2017) by plotting the fold-change of the average number of transposons per gene for libraries of each background, against a *p*-value associated with this difference (volcano plot, Fig. 3A). Clear differences stood out. First, a set of genes conferring arginine auxotrophy when mutated did not tolerate transposons in the W303 background. This can be explained by the fact that the W303 background bears a mutation in the arginine transporter *CAN1*, conferring Canavanine resistance to this strain (Thomas and Rothstein, 1989). As W303 cells are unable to import arginine, they are obligate arginine prototrophs. We found another striking difference in the genes encoding components of the RAM network (*TAO3*, *CBK1*, *KIC1*, *SOG2, MOB2* and *HYM1.* Note that *MOB2* scores low due to a large transposon-tolerant domain in all backgrounds) (Fig3A, B, C). This differential requirement for the RAM network in different laboratory strains was noted previously and attributed to a mutation in *SSD1* in the W303 background (Jorgensen et al., 2002), which renders the RAM network dispensable. Finally, we also found genes localized in the subtelomeric regions of the right arm of chromosome XIV, and in proximity of the YBRWTy1-2 repetitive element on chromosome II, regions that are entirely missing in the W303 background (Matheson et al., 2017).

**Fig. 3:**
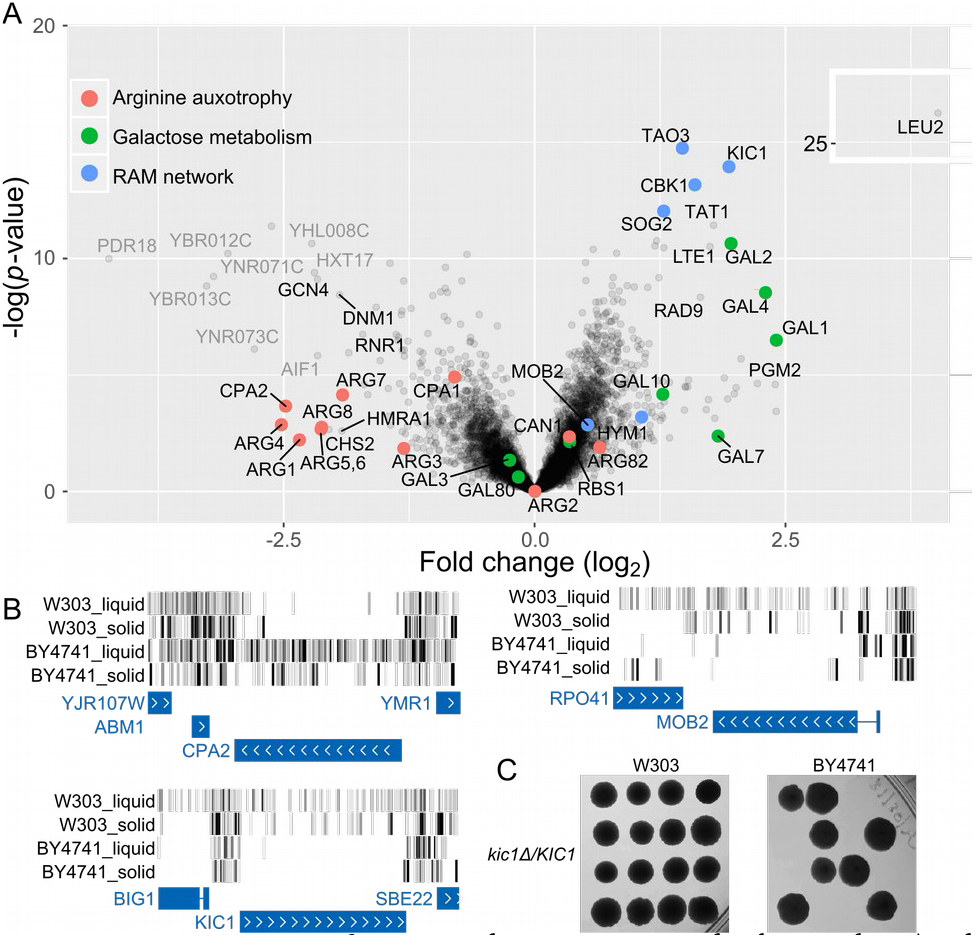
Comparison of W303 and By4741 genetic backgrounds. **A**) Volcano plot comparing the number of transposon insertions per gene between two W303-derived libraries and nine By4741-derived libraries. Genes with similar GO annotations are highlighted. Greyed-out genes are found on chromosomal regions entirely missing from the W303 background. **B**) Insertion maps of three genes annotated in (A), CPA2, MOB2 and KIC1. **C**) Assessment of the BY4741-specific lethality of kic1Δcells by tetrad dissection of indicating strains.

Consistent with the fact that the W303 and BY4741 backgrounds are closely related (Matheson et al., 2017), we found that their genetic requirement for growth were remarkably similar, while expected differences were standing out in our dataset. Therefore, our study lays the proof of principle that such inter-background comparisons, a topic of considerable interest (Hou et al., 2019), is easily feasible with the SATAY method.

### Comparing the Liquid- and Solid-Media Libraries

Comparing the two liquid libraries to the solid ones, we observed that the overall density of transposons per gene correlated well, and that most genes found to be essential by the solid approach were also essential in the liquid protocol. However, several genes that were intolerant to transposon insertions in the solid protocol, appeared unexpectedly studded with transposons in the liquid one. One class encompassed the genes for galactose utilization (*GAL1-11*, Fig. 4*A, B*). This stems from a fundamental difference between the solid and liquid approaches (Fig. 2A); in the solid procedure, strains are induced to transpose on +galactose -adenine plates. They must be able to form colonies on this medium and therefore require the *GAL* genes. In the liquid protocol, cells transpose in +galactose - uracil medium, but this happens in stationary phase, when the *GAL* genes are no longer necessary.

**Fig. 4:**
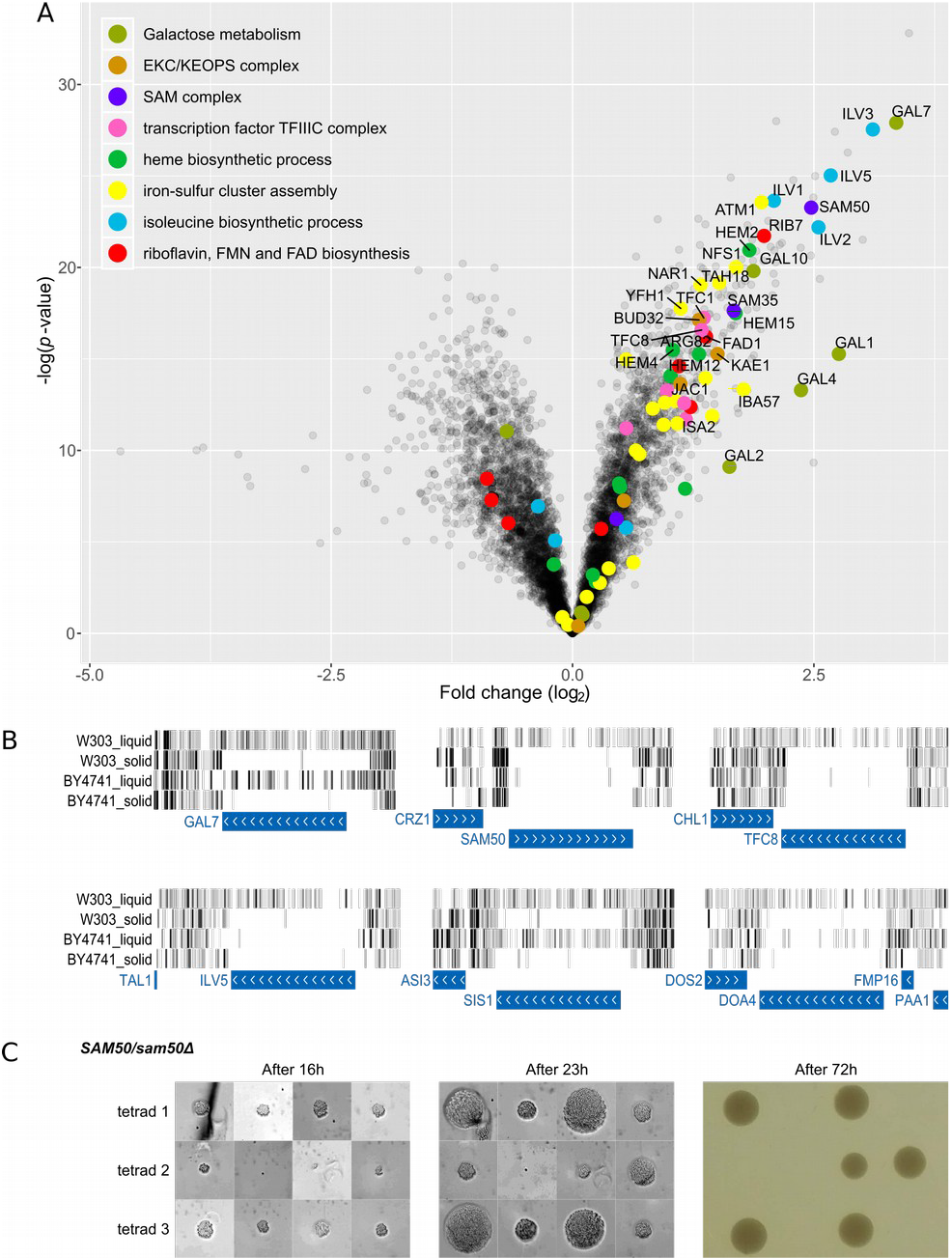
Comparison of libraries obtained following the liquid and solid protocols. **A**) Volcano plot comparing numbers of transposon insertions per gene between the liquid W303 library and 10 solid (one W303, and nine BY4741) libraries. Genes with indicated GO annotations are highlighted. **B**) Genomic maps showing transposon insertion data around interesting genes. **C**) A SAM50/sam50Δ heterozygous diploid was sporulated and tetrad dissected, and spore were allowed to grow for the indicated time (left and middle panel are microscopy images; right panel is a macroscopic observation).

More surprisingly, our analysis also revealed genes that encoded *bona fide* essential proteins (*Fig. 4A, B*). This was particularly clear in the liquid library generated in the W303 background. Striking examples were the components of the SAM complex *SAM50* and *SAM35*. The SAM complex is necessary for the proper insertion of the essential outer mitochondrial membrane protein translocon, Tom40 (Höhr et al., 2015). Therefore, finding multiple transposon insertions in both *SAM35* and *SAM50* but none in the *TOM40* gene was unintuitive. One other major difference between the liquid and solid library generation is timing; while in the solid protocol, each mutant has to form a micro-colony, which then needs be re-grown in selective medium, the liquid protocol skips the microcolony-forming step and drastically reduces the number of generations from the occurrence of the mutation to the final harvest of the library. Indeed, we could estimate that in the liquid protocol, only 10 (for the W303 library) to 14 (for the BY4741 one) generations elapsed from transposon insertion to harvest. We hypothesized that an essential gene might appear covered in transposons in liquid libraries if its deletion impairs growth with a delay. We tested the hypothesis that *sam50Δ* mutants could grow for a limited number of generations by creating *SAM50/sam50Δ* heterozygous diploids, sporulating them, and assessing the ability of *sam50Δ* spores to form micro-colonies (Fig. 4C). After 16h of growth, *sam50Δ* micro-colonies were indistinguishable from WT, and only later (23h) was the difference in growth apparent. Therefore, the liquid SATAY library generation protocol allows to detect essential genes with a phenotypic delay.

Other essential genes with phen0typic delays were genes involved in the biosynthesis of heme, riboflavin, iron-sulfur clusters, isoleucine and valine, as well as genes encoding components of the EKC/KEOPS and TFIIIC complexes (Fig. 4A, B).

While some phenotypically-delayed essential genes (like *SIS1*, Fig. 4B) were covered with transposons in both the BY4741 and the W303 liquid libraries, most (like *SAM50*, Fig. 4B) were only covered in the W303 liquid library, and many genes (like *DOA4*, Fig. 4B) had an intermediate transposon coverage in the BY4741 liquid library. These variations might be due to the different number of generations elapsed in the W303 (~10) and the BY4741 (~14) liquid libraries, and might pinpoint essential genes with phenotypic delays that manifest between 10 and 14 generations.

This is, to our knowledge, the first comprehensive dataset of essential genes with a phenotypic delay. A phenotypic delay might be expected if, for instance, the protein encoded or its product(s) is/are in vast excess over the amounts required for growth. In such case, the effect of a mutation will only be manifest after the factor has been diluted by a sufficient number of cell divisions. In principle, proteins in such excess could be found scattered in any pathway. Instead, our dataset highlights remarkably few well-defined pathways. Moreover, all the genes essential in those pathways exhibit a phenotypic delay.

This finding makes sense for essential protein complexes like SAM, TFIIIC or EKC/KEOPS, where all components are stoichiometrically expressed. It will be very interesting to determine what selective advantage justifies maintaining a select number of protein complexes at an apparently much higher concentration than needed.

Another class of phenotypically-delayed genes encodes near-complete biosynthesis pathways for metabolites (heme, Fe-S clusters, FAD, isoleucine and valine). Here, rather than the enzymes themselves, it is most likely the pathway’s end-product that is in large excess and responsible for the phenotypic delay. It is interesting that most of the molecules apparently biosynthesized in excess are redox cofactors. Though all of these cofactors play important roles in the electron transport chain, this is not what makes them essential in the conditions of the screen that use a fermentable carbon source (evidenced by the numerous transposon insertions within other respiratory genes). Fe-S clusters, for instance, are made in mitochondria but are essential components of several DNA metabolism enzymes in the nucleus, for reasons that are still mysterious. While heme and FAD are heavily involved in mitochondrial respiration, both are also essential for ergosterol biosynthesis, heme as part of the lanosterol 14α-demethylase and FAD as part of the Squalene epoxidase (Daum et al., 1998). Ergosterol biosynthesis is essential and the genes encoding the protein components of both enzymes, *ERG11* and *ERG1*, are devoid of transposons in all conditions. Additionally, FAD is necessary for oxidative protein folding (Tu and Weissman, 2002). Both cofactors thus appear to be synthesized in large excess over their need in ergosterol biosynthesis and thiol oxidation. The case of isoleucine and valine is intriguing. Although not mitochondrial redox cofactors, their biosynthesis happens within mitochondria and is connected to mitochondrial function. At least two enzymes of the pathway, Ilv5 and Ilv6, are components of the mitochondrial DNA nucleoid and are necessary for the maintenance of the mitochondrial genome. Why valine and isoleucine synthesis is connected to mitochondrial DNA is, to our knowledge, unclear.

Thus, while the timescale does not compare with the almost instantaneous inactivation that can be achieved, for instance, with Auxin-induced degrons (Nishimura et al., 2009), our liquid protocol allows assessing the consequences of mutations on a short timescale and detecting phenotypic delays at the genome scale. The number of generations can easily be controlled to assess the extent of the delay, which might hint on the fold excess of the protein/ product. The functional significance and the selective advantage of producing protein complexes and cofactors in apparent excess deserves further scrutiny.

### Screening for Drug Sensitivity and Resistance

With the new liquid protocol, a single investigator might easily generate dozens of SATAY libraries harboring gain-, loss-, and separation-of-function alleles. Loss-of-function mutations are generated by the disruption of a gene CDS, preventing the making of a functional protein. Gain-of-functions can be generated by the truncation of inhibitory domains, or by the insertion of a transposon in the gene promoter. This latter phenomenon happens because the transposon bears cryptic promoters (Yu et al., 2004) that can lead to the overexpression of downstream genes, as we have shown for *DDI1* (Serbyn et al., 2019). Separation-of-function alleles can be generated by selective truncation of functional domains, as exemplified by the TORC1 regulator *PIB2* (Michel et al., 2017). To assess the potential of our new protocol for generating such alleles, we created a liquid W303 library and challenged it with a compendium of 9 bioactive and antifungal drugs at sublethal concentrations. These drugs included inhibitors of lipid biosynthesis, glycosylation and cell wall biogenesis. After generation and re-growth of a transposon library containing an estimated 57 million independent clones, the library was split and grown for two successive rounds from OD=0.1 to saturation in the presence of compounds at an inhibitory concentration of 30% (IC_30_). This concentration allows identifying transposon insertions that confer either resistance or sensitivity to the compound (Hoepfner et al., 2014). Genomic DNA preparation was performed as previously described. All libraries were barcoded during amplification and simultaneously sequenced on a NextSeq platform.

The results of these screens are found in Supplementary Fig. 1 and 2 and supplementary dataset, and can be browsed here: http://genome-euro.ucsc.edu/s/benjou/2019compounds_r57ift.

Because we expected gain-of-functions to appear by transposon insertions in gene promoters (Serbyn et al., 2019), we not only analyzed the number of transposons and sequencing reads within CDS but also in promoter regions. Since there are no definitive ways to define promoter regions, we performed three analyses on (1) 200-bp upstream of the transcription start site as characterized by (Xu et al., 2009) (Supplementary Fig. 3), (2) 200-bp (Supplementary Fig. 4) and (3) 500-bp upstream of the ATG (Supplementary Fig. 5).

One straightforward way to cause resistance to an inhibitor of protein X is to overexpress protein X. Indeed, for the three compounds with characterized direct targets – Cerulenin, Fenpropimorph and Tunicamycin –, the promoters in which transposon insertions led to most resistance were precisely those of the direct targets of the drugs, i.e. *FAS1* and *ACC1* for Cerulenin (note that the unusually long 5’UTR of *FAS1* causes it to be missed when the promoter is defined relative to the ATG), *ERG24* and *ERG2* for Fenpropimorph, and *ALG7* for Tunicamycin (Fig. 5, top panel, Supplementary Fig. 3-5). Therefore, these screening conditions are ideally suited to discover direct targets of antifungal compounds.

**Fig. 5:**
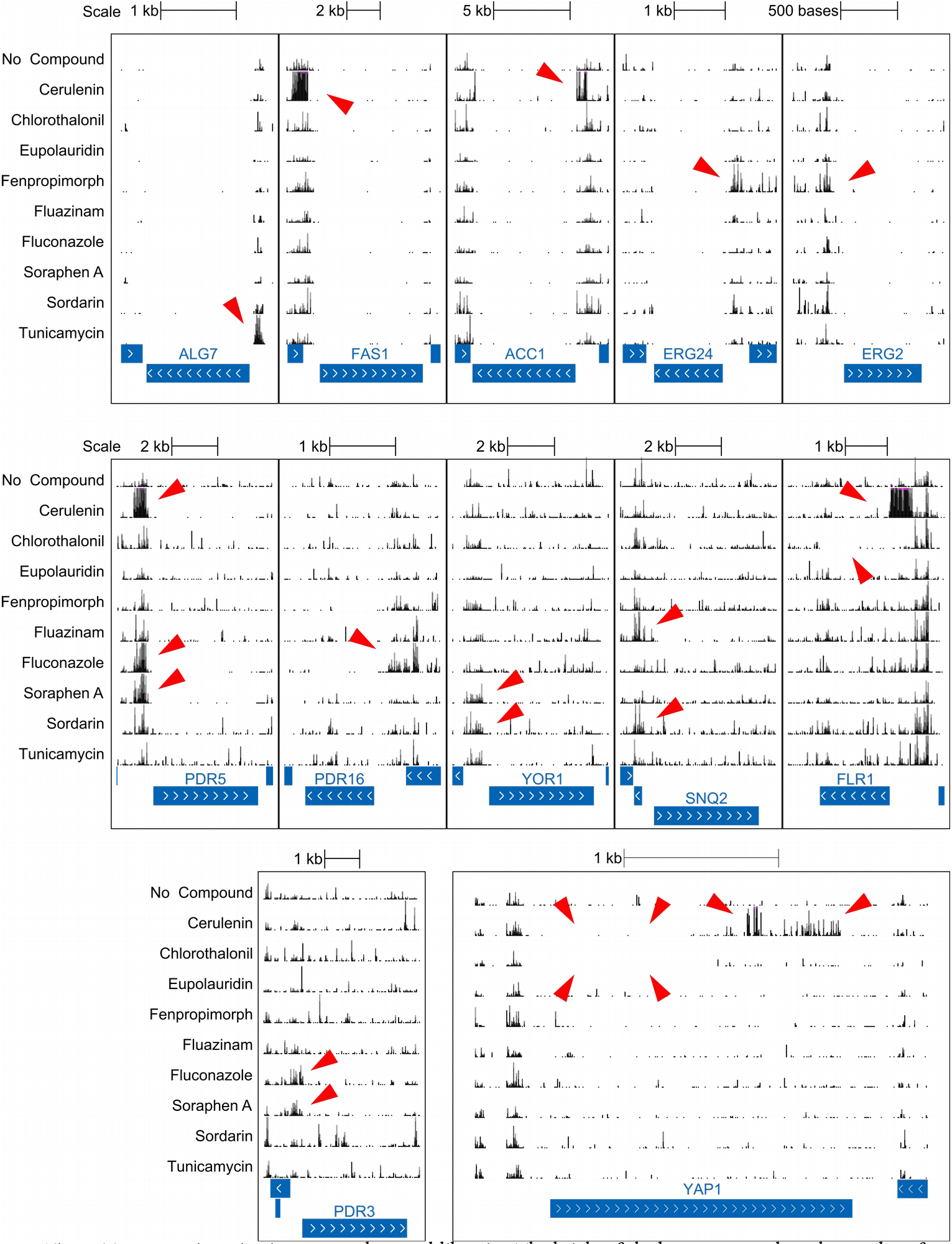
Transposon insertion in compound-treated libraries. The height of the bars corresponds to the number of sequencing reads, which reflects the fitness of the mutant. Top panel, red arrowheads indicate positively-selected insertions in the promoters of direct drug targets. Middle panel, red arrowheads indicate positively-selected insertions in the promoter of drug efflux transporters. Lower panel, red arrowhead indicate positively or negatively-selected insertions in the promoter and CDS of transcription factors PDR3 (master regulator of PDR5) and YAP1 (master regulator of FLR1).

Another way for a strain to become resistant to a drug is to overexpress a drug exporter. Indeed, we see numerous positively selected transposon insertions in the promoter of the multidrug efflux pump Pdr5 in libraries treated with Cerulenin, Fluconazole, Soraphen A and, to a lesser extent Sordarin (Fig. 5, middle panel, Supplementary Fig. 3-5). This was correlated with a positive selection for insertions in the promoter of *PDR3*, encoding a transcriptional activator of *PDR5* (Fig. 5, lower left panel, Supplementary Fig. 3-5). Other positively selected promoter insertions occurred for other multidrug transporters like *PDR16* in Fluconazole-treated, *YOR1* in Soraphen A-, Sordarin- and Tunicamycin-treated, *SNQ2* in Fluazinam- and Sordarin-treated, and *FLR1* in Cerulenin-treated libraries (Fig. 5, middle panel, Supplementary Fig. 3-5).

It is worth noting that while in several instances, positive selection for insertions in the promoter region correlated with negative selection for insertions in the gene CDS (e.g. *PDR5* in Cerulenin, Fluconazole and Soraphen A, Fig. 5, middle panel), this was not always the case, indicating that a gain- and a loss-of function did not necessarily yield opposite phenotypes. The case of *FLR1* is particularly interesting as loss of *FLR1* confers sensitivity to Chlorothalonil, while its overexpression likely confers resistance to Cerulenin (Fig. 5). This latter result suggests that Flr1 might be able to efficiently pump Cerulenin out of the cell. However, transposons in *FLR1* CDS are neither positively nor negatively selected in Cerulenin, suggesting that *FLR1* loss does not affect sensitivity to Cerulenin. A reasonable explanation for this conundrum is that *FLR1* might be poorly expressed in Cerulenin-treated conditions, in which case a loss-of-function might be of little consequence, while a gain-of-function in the form of a transcriptional boost might be highly beneficial. Indeed, *FLR1* is under the control of Yap1 (Oskouian and Saba, 1999), a redox-sensitive transcription factor. Yap1 is normally found in the cytoplasm. Oxidative stress causes oxidation and inactivation of a nuclear export sequence situated at the C-terminus of the protein, leading to nuclear import of the transcription factor and *FLR1* transcription induction (Gulshan et al., 2005; Kuge et al., 1997). We find that in the Cerulenin-treated library, insertions truncating the C-terminus, thus yielding a constitutively-active Yap1, are positively selected for, consistent with the idea that boosting *FLR1* expression is beneficial to respond to cerulenin (Fig. 5, lower right panel). Such transcriptional boost might not be needed in Chlorothalonil-treated cells, as Chlorothalonil wastes the pool of reduced gluthatione, hence creating oxidative conditions that likely cause oxidation of Yap1-nuclear export signal (Tillman et al., 1973). In support of this model, although Yap1 gain-of-function truncation does not confer any advantage in Chlorothalonil, loss-of-function mutations are lethal. A similar phenomenon was observed previously when we caused a synthetic growth defect between a *WSS1* deletion and an Auxin-induced degradation of Tdp1, indicating that Flr1 might pump out Auxin as well (Serbyn et al., 2019).

Thus, our data suggest that while *FLR1* is active and required for survival in Chlorothalonil, it is inactive in Cerulenin. Yet, a transcriptional boost, either by gain-of-function of Yap1 or direct mutation of *FLR1* promoter, is beneficial to survive exposure to a drug that Yap1 doesn’t normally respond to. Therefore, the evolutionary selection that keeps *FLR1* silent in non-stressed conditions might come with a cost; the inability to respond to drugs, like Cerulenin, that Yap1 cannot sense.

When we examine transposon insertions within CDSs, we can identify genes for which loss-of-function causes sensitivity or resistance to compounds. As seen previously, counting the number of transposons (Supplementary Fig. 1) and reads (Supplementary Fig. 2) per gene is better suited for assessing hypersensitivity and resistance, respectively. We observe, for instance, that loss of cryptic mating loci silencing via impairment of Sir2, Sir3 or Sir4, causes resistance to Eupolauridine, a potential DNA-damaging agent. This happens presumably because the loss of silencing causes the expression of both MATa and MATα information, triggering a “pseudodiploid” state, in which cells switch toward a homologous recombination-based DNA repair, which might repair the damages caused by Eupolauridine more efficiently.

Together these data show that SATAY is a valuable approach to identify (1) the direct targets of antifungal drugs, (2) the pumps responsible for drug excretion and whether these are active during drug treatment, and (3) mechanisms potentially leading to resistance.

### Conclusion

Our updated SATAY protocols allow the rapid and high throughput screening of loss-, gain- and separation-of-function mutations in yeast, and remove the burden of pouring, plating and harvesting hundreds of plates per screen. Transposon insertion sequencing is emerging as a method of choice to interrogate the genome of a growing number of micro-organisms. While *S. cerevisiae* has proven to be an outstanding tool for cell biology and biotechnology, deletion libraries are only available in select genetic backgrounds. We show that SATAY can be performed in different backgrounds and any condition with minimal manpower and material requirement. The vast scope of information that can be deducted from such screens is unfolding and we only present a cursory analysis of the dataset that we have generated. It is to be expected that with increasing number of screens, more sophisticated analysis tools, including machine-learning, will be highly successful to extract meaningful biological information from these data.

### Materials and Methods

#### Plasmids and yeast strains

Variants of the pBK257 plasmid with repair templates of various lengths and orientations were generated by PCR amplification of fragments of the *ADE2* gene (using primers #1 to #15, see table 1). The repair template was cloned by gap-repair in yeast after linearization of pBK257 with Kpn1. Yeast deletions were generated by PCR-mediated gene deletion using primers listed in table 1 using the Longtine and Janke toolboxes (Janke et al., 2004; Longtine et al., 1998).

**Table 1:**
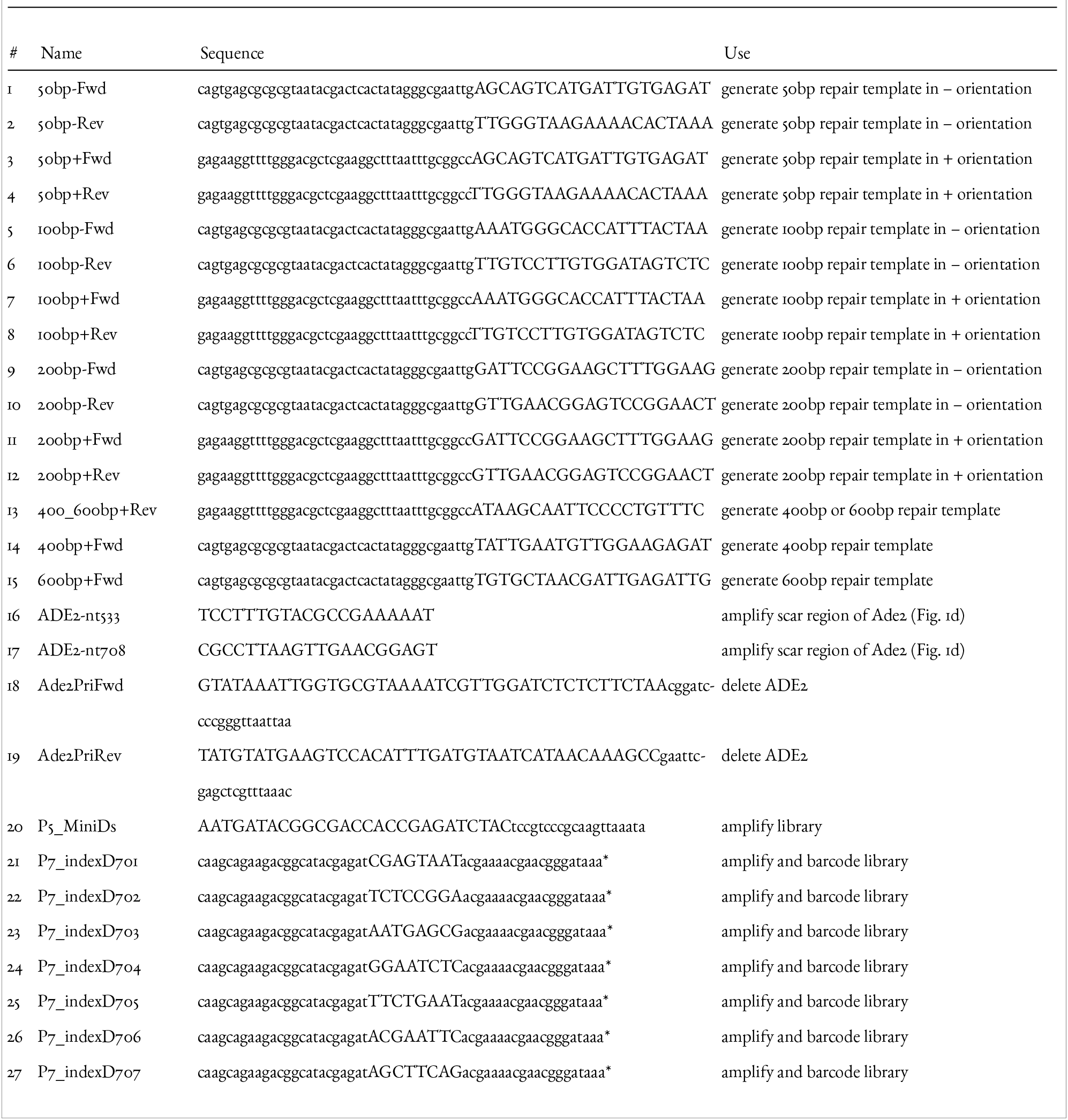
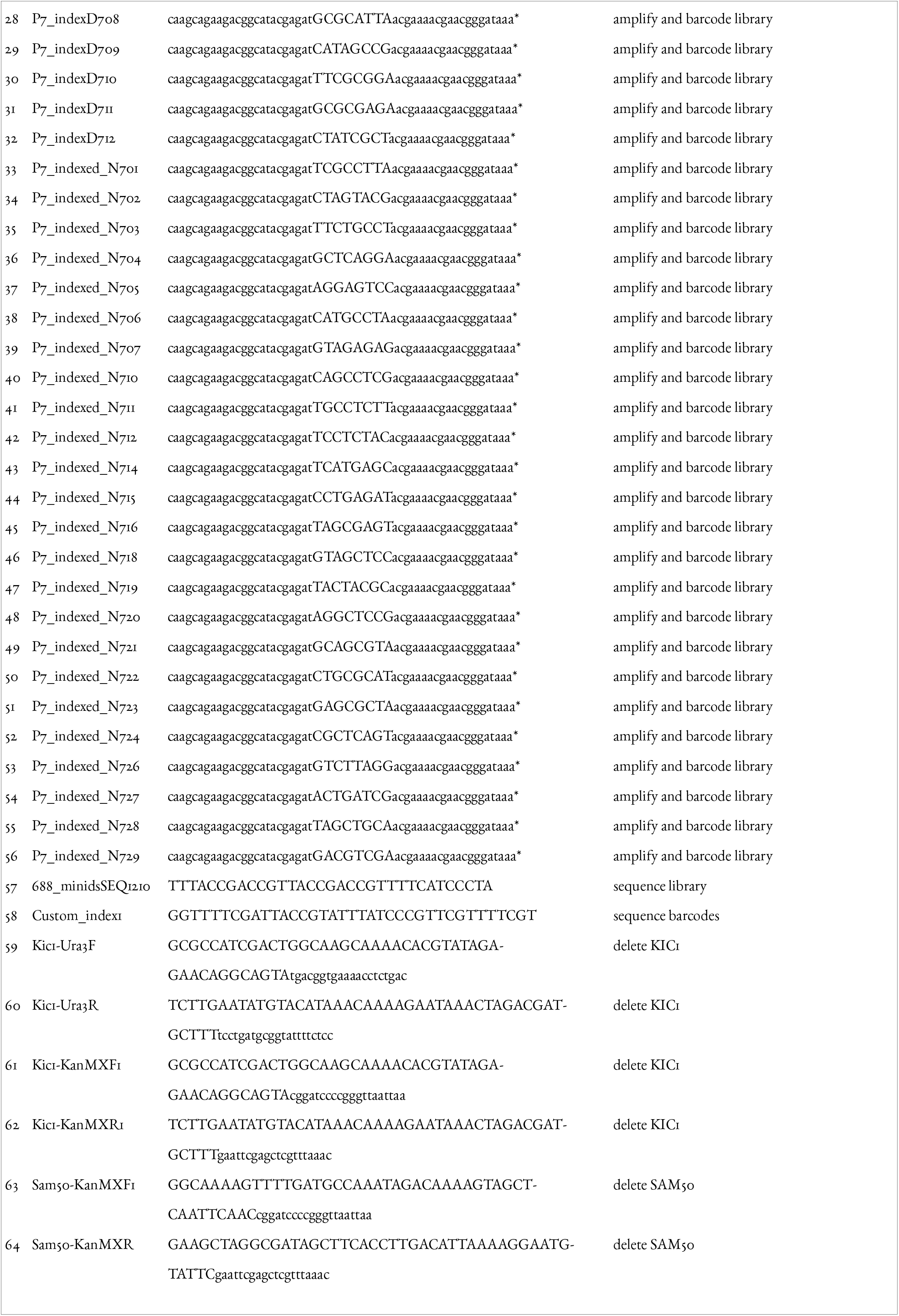
primers used in this study. *Oligonucleotide sequences © 2016 Illumina, Inc. All rights reserved. Derivative works created by Illumina customers are authorized for use with Illumina instruments and products only. All other uses are strictly prohibited

#### Libraries generation

yeast strains were transformed with pBK549. Individual clones were streaked on SC -URA and SC -ADE plates. Typically three to six clones that produced either no or just a few colonies on SD -ADE were used further. The SC -URA plates were used to inoculate a pre-culture in SC -URA + 2%Raffinose +0.2% Glucose at an OD of 0.2, then grown for 16hours to saturation.

For solid libraries, the saturated pre-culture was concentrated to OD39. 200microl of the concentrated culture were plated on 50 to 100 SC -ADE + 2%Galactose plates. Cells were harvested after 15-21 days and processed as described previously (Michel et al. 2017).

For liquid libraries, saturated pre-cultures were diluted to OD 0.2 in SC -URA + 2%Galactose medium and grown for 51-57 hours. Saturated induced cultures were then diluted into SC -ADE + 2%Glucose and grown for ~70h to OD ~2. Cells were harvested and processed for DNA extraction and sequencing. 200microl of the galactose induced culture or dilutions of it were plated at t_0_, t_20_ and at the end of induction on SC, SC -URA and SC - ADE plates. Platings at t_0_ served to estimate the background of ADE+ clones due to spontaneous recombination of pBK549 (typically 0.01% in By4741 background and 0.007% in W303 background). Platings at t_20_ were counted at the end of the induction to guide the re-inoculation following transposition. The number of ADE+ cells at t_20_ typically corresponds to ~10% of the number of ADE+ cells at the end of induction.

#### DNA extraction and preparation

DNA was extracted from 0.5 g yeast pellets as described previously (Michel et al., 2017), and processed by restriction digestion, circularization and PCR amplification. Barcodes were introduced in the PCR product by the use of primer #20 in combination with either of primers #21 to #56. Library chaaracteristics are described in Table 4.

**Table 2:** Yeast strains used in this study

| Name | Parent | Genotype | Reference |
| --- | --- | --- | --- |
| ByK352 | BY4741 | MATa his3Δ1 leu2Δo met17Δo ura3Δo ade2Δ::HIS3 | Michel et al. 2017 |
| ByK485 | ByK4741/<br>ByK4742 | MATa/α his3Δ1/his3Δ1 leu2Δo/leu2Δo LYS2/lys2Δo met17Δo/<br>MET17 ura3Δo/ura3Δo ade2Δ::HIS3/ade2Δ::HIS3 | Michel et al. 2017 |
|  | ByK485 | MATa/α his3Δ1/his3Δ1 leu2Δo/leu2Δo LYS2/lys2Δo met17Δo/<br>MET17 ura3Δo/ura3Δo ade2Δ::HIS3/ade2Δ::HIS3 SAM50/<br>Sam50::URA3 | This study |
| ByK831 | W303-1A | MATa {leu2-3,112 trp1-1 can1-100 ura3-1 ade2-1 his3-11,15} |  |
| ByK912 | W303-1A | MATa {leu2-3,112 trp1-1 can1-100 ura3-1 ade2-1 his3-11,15} |  |
| ByK832 | ByK831 | MATα {leu2-3,112 trp1-1 can1-100 ura3-1 ade2-1 his3-11,15}<br>ade2Δ::NatMx6(?)* | This study |
| ByK830 | W303-1B | MATα {leu2-3,112 trp1-1 can1-100 ura3-1 ade2-1 his3-11,15} |  |
| ByK911 | W303-1B | MATalpha {leu2-3,112 trp1-1 can1-100 ura3-1 ade2-1 his3-11,15} |  |
| ByK999 | ByK911/<br>ByK912 | MATa/MATα {leu2-3,112/leu2-3,112 trp1-1/trp1-1 can1-100/can1-100<br>ura3-1/ ura3-1 ade2-1/ ade2-1 his3-11,15/his3-11,15} KIC1/<br>Kic1::KanMX6 | This study |
| ByK1000 | ByK911/<br>ByK912 | MATa/MATα {leu2-3,112/leu2-3,112 trp1-1/trp1-1 can1-100/can1-100<br>ura3-1/ ura3-1 ade2-1/ ade2-1 his3-11,15/his3-11,15} SAM50/<br>Sam50::KanMX6 | This study |

**Table 3:**
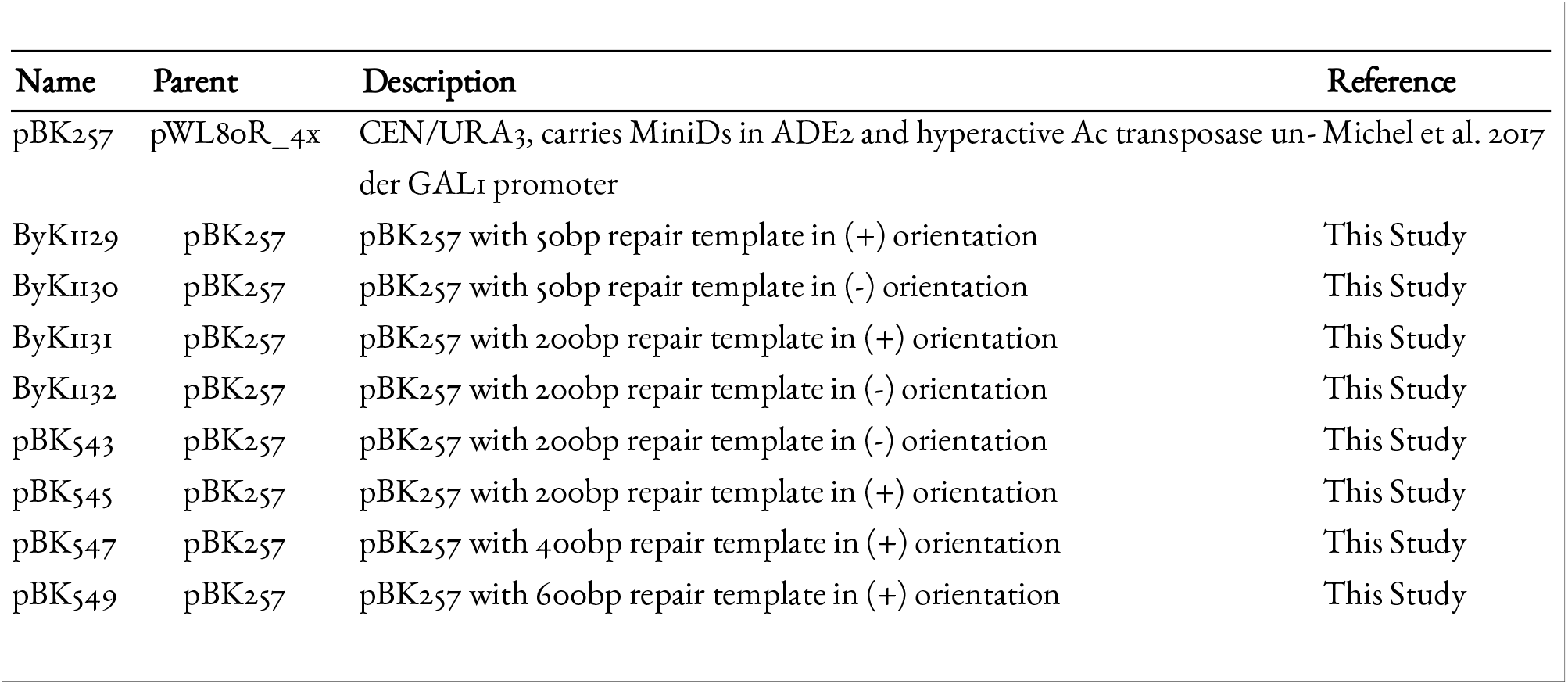
Plasmids used in this study

| Name | Parent | Description | Reference |
| --- | --- | --- | --- |
| pBK257 | pWL8oR_4x | CEN/URA3, carries MiniDs in ADE2 and hyperactive Ac transposase under GAL1 promoter | Michel et al. 2017 |
| ByK1129 | pBK257 | pBK257 with 50bp repair template in (+) orientation | This Study |
| ByK1130 | pBK257 | pBK257 with 50bp repair template in (-) orientation | This Study |
| ByK1131 | pBK257 | pBK257 with 200bp repair template in (+) orientation | This Study |
| ByK1132 | pBK257 | pBK257 with 200bp repair template in (-) orientation | This Study |
| pBK543 | pBK257 | pBK257 with 200bp repair template in (-) orientation | This Study |
| pBK545 | pBK257 | pBK257 with 200bp repair template in (+) orientation | This Study |
| pBK547 | pBK257 | pBK257 with 400bp repair template in (+) orientation | This Study |
| pBK549 | pBK257 | pBK257 with 600bp repair template in (+) orientation | This Study |

**Table 4:** Characteristics of the libraries presented in this study

| library | Total number of clones (sub-libraries) | transposons mapped | reads mapped | mean reads per transposon | median reads per transposon |
| --- | --- | --- | --- | --- | --- |
| BY4741 liquid | $\sim 3.8 \cdot 10^6$ (11.4+10.4+10.1+6.7)·10<sup>5</sup> | 371626 | 16398184 | 44.13 | 11 |
| BY4741 solid | $\sim 3.4 \cdot 10^6$ (8.6+7.0+9+9.3)·10<sup>4</sup> | 310566 | 9548788 | 30.75 | 6 |
| W303 Liquid | $\sim 12 \cdot 10^6$ | 708320 | 16604290 | 23.44 | 5 |
| W303 Solid | $\sim 3.4 \cdot 10^6$ (1.96+1.46)·10<sup>6</sup> | 401635 | 18282456 | 45.52 | 9 |
| Sordarin | $\sim 56 \cdot 10^6$ (14.6+12.8+16.4+13)·10<sup>6</sup> | 340906 | 9328340 | 27.36 | 5 |
| Tunicamycin | $\sim 56 \cdot 10^6$ (14.6+12.8+16.4+13)·10<sup>6</sup> | 343104 | 11033972 | 32.16 | 6 |
| Cerulenin | $\sim 56 \cdot 10^6$ (14.6+12.8+16.4+13)·10<sup>6</sup> | 340870 | 9171326 | 26.91 | 5 |
| Fluconazole | $\sim 56 \cdot 10^6$ (14.6+12.8+16.4+13)·10<sup>6</sup> | 329616 | 8029800 | 24.36 | 5 |
| Fenpropimorph | $\sim 56 \cdot 10^6$ (14.6+12.8+16.4+13)·10<sup>6</sup> | 338561 | 8506055 | 25.12 | 5 |
| Soraphen A | $\sim 56 \cdot 10^6$ (14.6+12.8+16.4+13)·10<sup>6</sup> | 349968 | 5960216 | 17.03 | 5 |
| Eupolauridine | $\sim 56 \cdot 10^6$ (14.6+12.8+16.4+13)·10<sup>6</sup> | 275227 | 4567823 | 16.6 | 4 |
| Fluazinam | $\sim 56 \cdot 10^6$ (14.6+12.8+16.4+13)·10<sup>6</sup> | 311038 | 7722266 | 24.83 | 5 |
| Chlorothalonil | $\sim 56 \cdot 10^6$ (14.6+12.8+16.4+13)·10<sup>6</sup> | 282976 | 7932777 | 28.03 | 6 |
| No Compound | $\sim 56 \cdot 10^6$ (14.6+12.8+16.4+13)·10<sup>6</sup> | 341674 | 7550413 | 22.1 | 5 |

#### Compound treatments

the following compounds were resuspended in DMSO as a 1000x stock. A freshly induced liquid yeast tranposon library was diluted to OD 0.2 into SC -ADE + 2%Glucose and grown to OD 1. It was then diluted to OD 0.1 in 500 ml of the same medium, with the following compound concentrations: Cerulenin, 0.11 mg/l; Chlorothalonil, 0.086 mg/l; Eupolauridine, 4.18 mg/l; Fenpropimorph, 1.7 mg/l; Fluazinam, 0.047 mg/l; Fluconazole, 5.68 mg/l; Soraphen A, 0.048 mg/ml; Sordarin, 0.085 mg/ml, Tunicamycin, 0.22 mg/ml. Cultures were grown to saturation and re-diluted in 500 ml for a repeated treatment with or without compound from OD 0.1 to saturation.

## Supplementary Figure Legends

**Supplementary Fig. 1:**
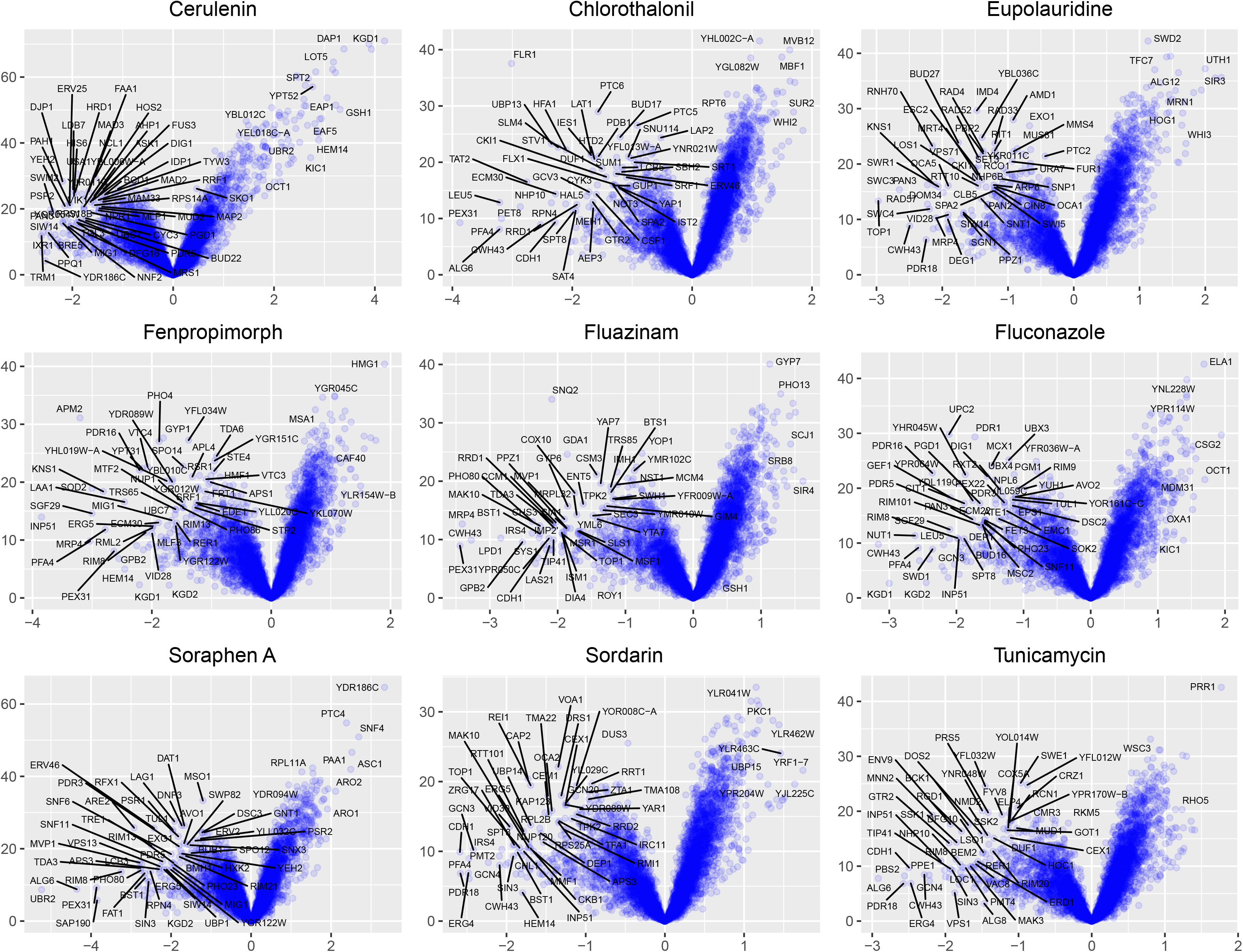
Volcano plot analysis for the number of transposons mapping in the CDS of genes for each of the compounds.

**Supplementary Fig. 2:**
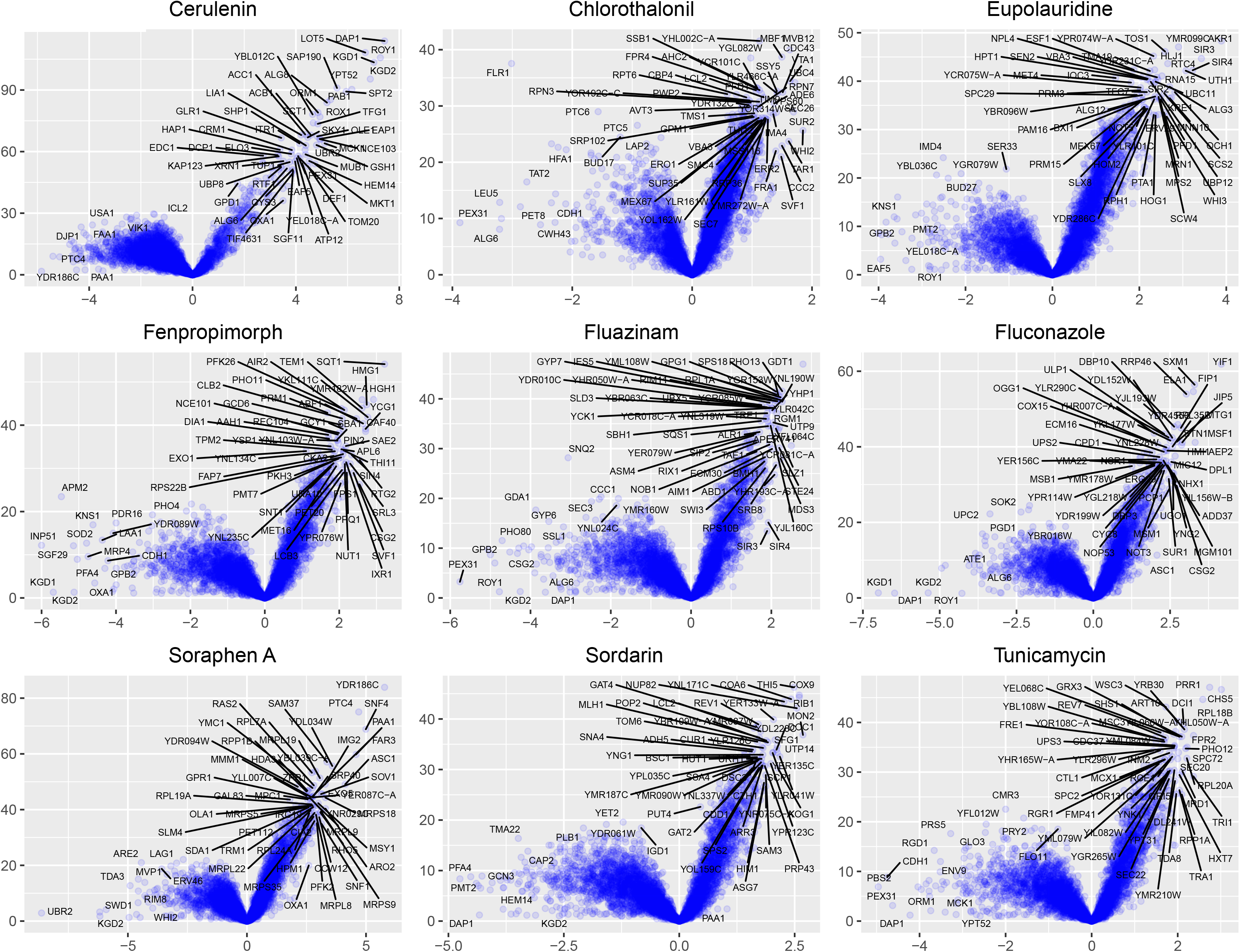
Volcano plot analysis for the number of sequencing reads mapping in the CDS of genes for each of the compounds.

**Supplementary Fig. 3:**
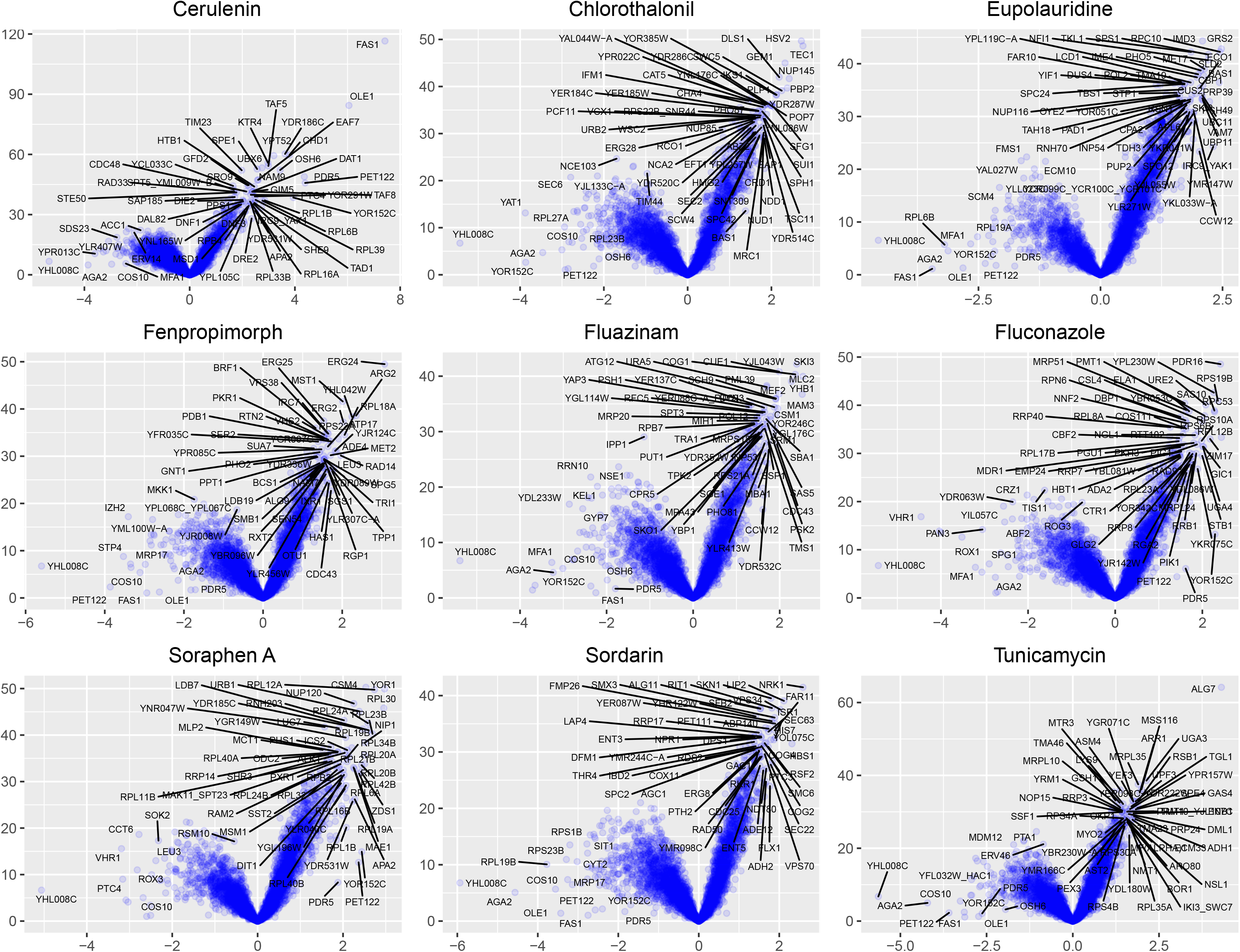
Volcano plot analysis for the number of sequencing reads mapping in the promoters of genes for each of the compounds. Promoters are defined as the 200 bp upstream of the transcription start sites as assessed by Xu et al.

**Supplementary Fig. 4:**
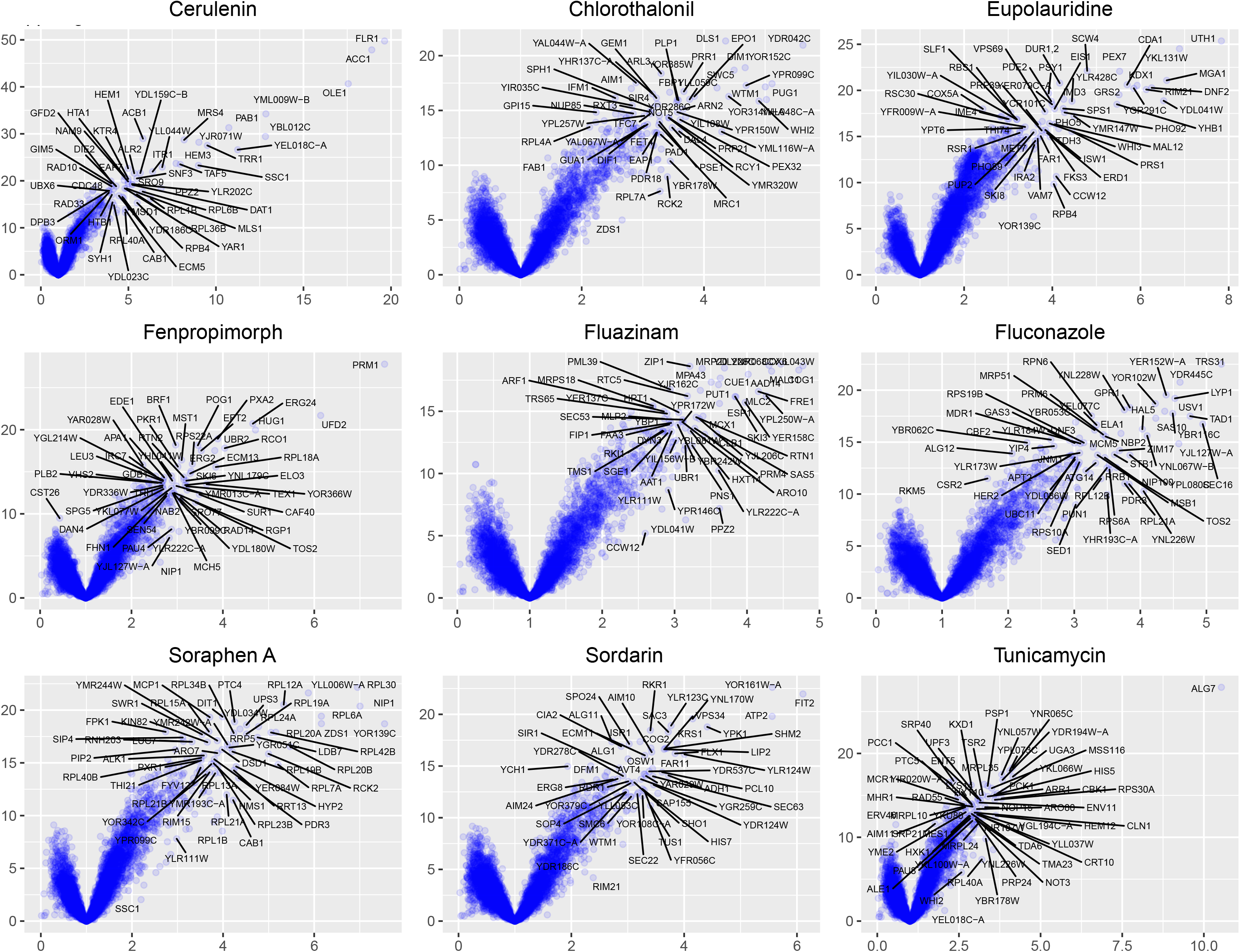
Volcano plot analysis for the number of sequencing reads mapping in the promoters of genes for each of the compounds. Promoters are defined as the 200 bp upstream of the translation start site.

**Supplementary Fig. 5:**
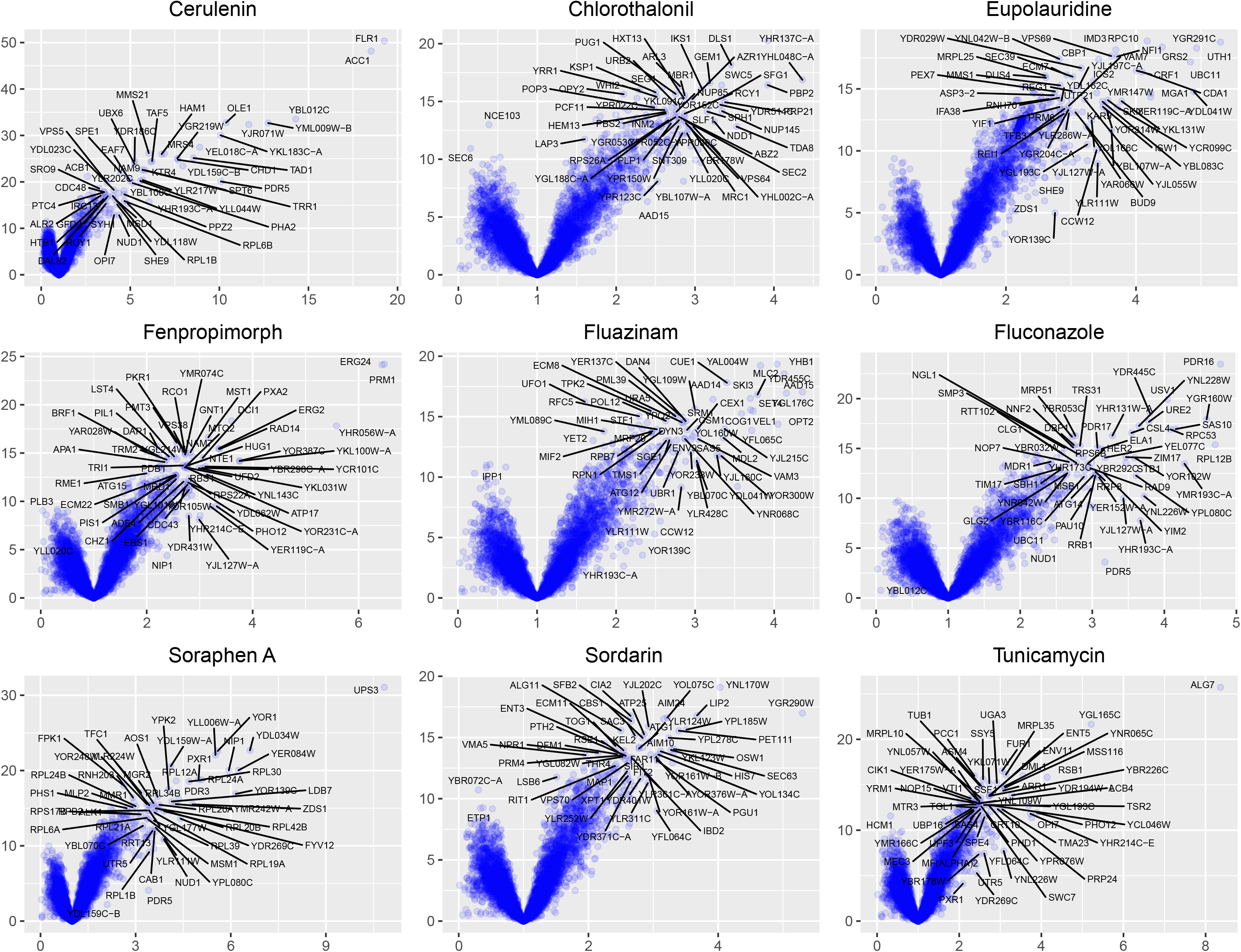
Volcano plot analysis for the number of sequencing reads mapping in the promoters of genes for each of the compounds. Promoters are defined as the 500 bp upstream of the translation start site.

## References

Boulton SJ, Jackson SP. 1996. Identification of a Saccharomyces cerevisiae Ku80 homologue: roles in DNA double strand break rejoining and in telomeric maintenance. Nucleic Acids Res 24:4639–4648.

Christen B, Abeliuk E, Collier JM, Kalogeraki VS, Passarelli B, Coller JA, Fero MJ, McAdams HH, Shapiro L. 2014. The essential genome of a bacterium. Molecular Systems Biology 7:528–528. doi:10.1038/msb.2011.58

Coradetti ST, Pinel D, Geiselman GM, Ito M, Mondo SJ, Reilly MC, Cheng Y-F, Bauer S, Grigoriev IV, Gladden JM, Simmons BA, Brem RB, Arkin AP, Skerker JM. 2018. Functional genomics of lipid metabolism in the oleaginous yeast Rhodosporidium toruloides. Elife 7. doi:10.7554/eLife.32110

Daum G, Lees ND, Bard M, Dickson R. 1998. Biochemistry, cell biology and molecular biology of lipids of Saccharomyces cerevisiae. Yeast 14:1471–1510. doi:10.1002/(SICI)1097-0061(199812)14:16<1471::AID-YEA353>3.0.CO;2-Y

Edskes HK, Mukhamedova M, Edskes BK, Wickner RB. 2018. Hermes Transposon Mutagenesis Shows [URE3] Prion Pathology Prevented by a Ubiquitin-Targeting Protein: Evidence for Carbon/Nitrogen Assimilation Cross Talk and a Second Function for Ure2p in Saccharomyces cerevisiae. Genetics 209:789–800.

Giaever G, Chu AM, Ni L, Connelly C, Riles L, Véronneau S, Dow S, Lucau-Danila A, Anderson K, André B, Arkin AP, Astromoff A, El-Bakkoury M, Bangham R, Benito R, Brachat S, Campanaro S, Curtiss M, Davis K, Deutschbauer A, Entian K-D, Flaherty P, Foury F, Garfinkel DJ, Gerstein M, Gotte D, Güldener U, Hegemann JH, Hempel S, Herman Z, Jaramillo DF, Kelly DE, Kelly SL, Kötter P, LaBonte D, Lamb DC, Lan N, Liang H, Liao H, Liu L, Luo C, Lussier M, Mao R, Menard P, Ooi SL, Revuelta JL, Roberts CJ, Rose M, Ross-Macdonald P, Scherens B, Schimmack G, Shafer B, Shoemaker DD, Sookhai-Mahadeo S, Storms RK, Strathern JN, Valle G, Voet M, Volckaert G, Wang C, Ward TR, Wilhelmy J, Winzeler EA, Yang Y, Yen G, Youngman E, Yu K, Bussey H, Boeke JD, Snyder M, Philippsen P, Davis RW, Johnston M. 2002. Functional profiling of the Saccharomyces cerevisiae genome. Nature 418:387–391. doi:10.1038/nature00935

Giaever G, Nislow C. 2014. The yeast deletion collection: a decade of functional genomics. Genetics 197:451–465. doi:10.1534/genetics.114.161620

Girgis HS, Liu Y, Ryu WS, Tavazoie S. 2007. A Comprehensive Genetic Characterization of Bacterial Motility. PLOS Genetics 3:e154. doi:10.1371/journal.pgen.0030154

Gulshan K, Rovinsky SA, Coleman ST, Moye-Rowley WS. 2005. Oxidant-specific Folding of Yap1p Regulates Both Transcriptional Activation and Nuclear Localization. J Biol Chem 280:40524–40533. doi:10.1074/jbc.M504716200

Guo Y, Park JM, Cui B, Humes E, Gangadharan S, Hung S, FitzGerald PC, Hoe K-L, Grewal SIS, Craig NL, Levin HL. 2013. Integration Profiling of Gene Function With Dense Maps of Transposon Integration. Genetics 195:599–609. doi:10.1534/genetics.113.152744

Hoepfner D, Helliwell SB, Sadlish H, Schuierer S, Filipuzzi I, Brachat S, Bhullar B, Plikat U, Abraham Y, Altorfer M, Aust T, Baeriswyl L, Cerino R, Chang L, Estoppey D, Eichenberger J, Frederiksen M, Hartmann N, Hohendahl A, Knapp B, Krastel P, Melin N, Nigsch F, Oakeley EJ, Petitjean V, Petersen F, Riedl R, Schmitt EK, Staedtler F, Studer C, Tallarico JA, Wetzel S, Fishman MC, Porter JA, Movva NR. 2014. High-resolution chemical dissection of a model eukaryote reveals targets, pathways and gene functions. Microbiological Research 169:107–120. doi:10.1016/j.micres.2013.11.004

Höhr AIC, Straub SP, Warscheid B, Becker T, Wiedemann N. 2015. Assembly of β-barrel proteins in the mitochondrial outer membrane. Biochim Biophys Acta 1853:74–88. doi:10.1016/j.bbamcr.2014.10.006

Hou J, Tan G, Fink GR, Andrews BJ, Boone C. 2019. Complex modifier landscape underlying genetic background effects. PNAS 116:5045–5054. doi:10.1073/pnas.1820915116

Hua S, Qiu M, Chan E, Zhu L, Luo Y. 1997. Minimum Length of Sequence Homology Required forin VivoCloning by Homologous Recombination in Yeast. Plasmid 38:91–96. doi:10.1006/plas.1997.1305

Janke C, Magiera MM, Rathfelder N, Taxis C, Reber S, Maekawa H, Moreno-Borchart A, Doenges G, Schwob E, Schiebel E, Knop M. 2004. A versatile toolbox for PCR-based tagging of yeast genes: new fluorescent proteins, more markers and promoter substitution cassettes. Yeast (Chichester, England) 21:947–62. doi:10.1002/yea.1142

Jorgensen P, Nelson B, Robinson MD, Chen Y, Andrews B, Tyers M, Boone C. 2002. High-resolution genetic mapping with ordered arrays of Saccharomyces cerevisiae deletion mutants. Genetics 162:1091–1099.

Kuge S, Jones N, Nomoto A. 1997. Regulation of yAP-1 nuclear localization in response to oxidative stress. The EMBO Journal 16:1710–1720. doi:10.1093/emboj/16.7.1710

Lazarow K, Du M-L, Weimer R, Kunze R. 2012. A Hyperactive Transposase of the Maize Transposable Element Activator (Ac). Genetics 191:747–756. doi:10.1534/genetics.112.139642

Longtine MS, McKenzie A, Demarini DJ, Shah NG, Wach A, Brachat A, Philippsen P, Pringle JR. 1998. Additional modules for versatile and economical PCR-based gene deletion and modification in Saccharomyces cerevisiae. Yeast (Chichester, England) 14:953–61. doi:10.1002/(SICI)1097-0061(199807)14:10<953::AID-YEA293>3.0.CO;2-U

Matheson K, Parsons L, Gammie A. 2017. Whole-Genome Sequence and Variant Analysis of W303, a Widely-Used Strain of Saccharomyces cerevisiae. G3 (Bethesda) 7:2219–2226. doi:10.1534/g3.117.040022

Michel AH, Hatakeyama R, Kimmig P, Arter M, Peter M, Matos J, De Virgilio C, Kornmann B. 2017. Functional mapping of yeast genomes by saturated transposition. eLife 6. doi:10.7554/eLife.23570

Nishimura K, Fukagawa T, Takisawa H, Kakimoto T, Kanemaki M. 2009. An auxin-based degron system for the rapid depletion of proteins in nonplant cells. Nature Methods 6:917–922. doi:10.1038/nmeth.1401

Oskouian B, Saba JD. 1999. YAP1 confers resistance to the fatty acid synthase inhibitor cerulenin through the transporter Flr1p in Saccharomyces cerevisiae. Mol Gen Genet 261:346–353. doi:10.1007/s004380050975

Sanchez MR, Payen C, Cheong F, Hovde BT, Bissonnette S, Arkin AP, Skerker JM, Brem RB, Caudy AA, Dunham MJ. 2019. Transposon insertional mutagenesis in Saccharomyces uvarum reveals trans-acting effects influencing species-dependent essential genes. Genome Res 29:396–406. doi:10.1101/gr.232330.117

Segal ES, Gritsenko V, Levitan A, Yadav B, Dror N, Steenwyk JL, Silberberg Y, Mielich K, Rokas A, Gow NAR, Kunze R, Sharan R, Berman J. 2018. Gene Essentiality Analyzed by In Vivo Transposon Mutagenesis and Machine Learning in a Stable Haploid Isolate of Candida albicans. mBio 9:e02048–18. doi:10.1128/mBio.02048-18

Serbyn N, Noireterre A, Bagdiul I, Plank M, Michel AH, Loewith R, Kornmann B, Stutz F. 2019. The Aspartic Protease Ddi1 Contributes to DNA-Protein Crosslink Repair in Yeast. bioRxiv 575860. doi:10.1101/575860

Teng X, Dayhoff-Brannigan M, Cheng W-C, Gilbert CE, Sing CN, Diny NL, Wheelan SJ, Dunham MJ, Boeke JD, Pineda FJ, Hardwick JM. 2013. Genome-wide Consequences of Deleting Any Single Gene. Molecular Cell 52:485–494. doi:10.1016/j.molcel.2013.09.026

Thomas BJ, Rothstein R. 1989. Elevated recombination rates in transcriptionally active DNA. Cell 56:619–630. doi:10.1016/0092-8674(89)90584-9

Tillman RW, Siegel MR, Long JW. 1973. Mechanism of action and fate of the fungicide chlorothalonil (2,4,5,6-tetrachloroisophthalonitrile) in biological systems: I. Reactions with cells and subcellular components of Saccharomyces pastorianus. Pesticide Biochemistry and Physiology 3:160–167. doi:10.1016/0048-3575(73)90100-4

Tu BP, Weissman JS. 2002. The FAD- and O(2)-dependent reaction cycle of Ero1-mediated oxidative protein folding in the endoplasmic reticulum. Mol Cell 10:983–994.

Uhse S, Pflug FG, Stirnberg A, Ehrlinger K, Haeseler A von, Djamei A. 2018. In vivo insertion pool sequencing identifies virulence factors in a complex fungal–host interaction. PLOS Biology 16:e2005129. doi:10.1371/journal.pbio.2005129

van Opijnen T, Bodi KL, Camilli A. 2009. Tn-seq: high-throughput parallel sequencing for fitness and genetic interaction studies in microorganisms. Nat Methods 6:767–772. doi:10.1038/nmeth.1377

Weil CF, Kunze R. 2000. Transposition of maize Ac/Ds transposable elements in the yeast Saccharomyces cerevisiae. Nat Genet 26:187–190. doi:10.1038/82827

Weissman J, Guthrie C, Fink GR. 2010. Guide to Yeast Genetics: Functional Genomics, Proteomics, and Other Systems Analysis. Academic Press.

Xu Z, Wei W, Gagneur J, Perocchi F, Clauder-Münster S, Camblong J, Guffanti E, Stutz F, Huber W, Steinmetz LM. 2009. Bidirectional promoters generate pervasive transcription in yeast. Nature 457:1033–1037. doi:10.1038/nature07728

Yu J, Marshall K, Yamaguchi M, Haber JE, Weil CF. 2004. Microhomology-Dependent End Joining and Repair of Transposon-Induced DNA Hairpins by Host Factors in Saccharomyces cerevisiae. Molecular and Cellular Biology 24:1351–1364. doi:10.1128/MCB.24.3.1351-1364.2004

Zhu J, Gong R, Zhu Q, He Q, Xu N, Xu Y, Cai M, Zhou X, Zhang Y, Zhou M. 2018. Genome-Wide Determination of Gene Essentiality by Transposon Insertion Sequencing in Yeast Pichia pastoris. Scientific Reports 8:10223. doi:10.1038/s41598-018-28217-z

